# Differential retention of transposable element-derived sequences in outcrossing Arabidopsis genomes

**DOI:** 10.1101/368985

**Authors:** Sylvain Legrand, Thibault Caron, Florian Maumus, Sol Schvartzman, Leandro Quadrana, Eléonore Durand, Sophie Gallina, Maxime Pauwels, Clément Mazoyer, Lucie Huyghe, Vincent Colot, Marc Hanikenne, Vincent Castric

## Abstract

Transposable elements (TEs) have initially been viewed as pure genomic parasites but are now recognized as central genome architects and powerful agents of rapid adaptation. A proper evaluation of their evolutionary significance has been hampered by the paucity of short scale phylogenetic comparisons between closely related species. Here, we characterized the dynamics of TE accumulation at the micro-evolutionary scale by comparing two closely related plant species, *Arabidopsis lyrata* and *A. halleri*. Joint genome annotation in these two outcrossing species confirmed that both contain two distinct populations of TEs with either ‘recent’ or ‘old’ insertion histories. Identification of rare segregating insertions suggests that diverse TE families contribute to the ongoing dynamics of TE accumulation in the two species. TE fragments that have been maintained in both species, *i.e.* those that are orthologous, tend to be located on average closer to genes than those that are retained in one species only. Moreover, compared to non-orthologous TE insertions, those that are orthologous tend to produce fewer short interfering RNAs, are less heavily methylated when found within or adjacent to genes and these tend to have lower expression levels. These findings suggest that long-term retention of TE insertions reflects their frequent acquisition of adaptive roles and/or the deleterious effects of removing TE insertions when they are close to genes. Overall, our results indicate a rapid evolutionary dynamics of the TE landscape in these two outcrossing species, with an important input of a diverse set of new insertions with variable propensity to resist deletion.

## INTRODUCTION

Transposable elements (TEs) are repeated elements found almost universally in eukaryotic genomes that can proliferate by high-jacking a variety of cellular processes. They are believed to be the substrate over which the non-coding fraction of the genome is formed in the long term (Britten and Kohne 1968) and contribute a large fraction of genome size variation across taxa, representing as much as 85% of the maize and barley genome and around 20% in *A. thaliana* (Arabidopsis Genome Initiative 2000; Wicker et al. 2005; Buisine et al. 2008; Schnable et al. 2009). Their spread in genomes is limited by mechanisms to suppress their transposition activity by host defense mechanisms including the production of dedicated classes of small non-coding RNAs (piRNA and siRNA) causing transcriptional silencing by RNA-dependent DNA methylation (RdDM) (Slotkin and Martienssen 2007).

In spite of their quantitative importance, the evolutionary significance of TEs has been the subject of constant debate in the field. Their discovery was immediately followed by the interpretation that they must represent important “controlling elements” (McClintock 1950) that confer selective advantages to the organism and are a major “fuel” for evolution (Slotkin and Martienssen 2007; Rebollo et al. 2012). This interpretation was soon challenged by the realization that TEs propagate in a largely selfish manner, and a large body of literature has considered them essentially as genomic parasites (Orgel and Crick 1980). Over the last decade, however, molecular studies have reported convincing examples of TEs determining important evolutionary novelties and contributing to essential biological functions such as the rewiring of entire transcriptional networks along sex chromosomes (Ellison and Bachtrog 2015). Several iconic examples of rapid adaptive evolution have been linked to TE insertions such as the industrial melanism in the peppered moth (Van’t Hof et al. 2016) or the change in branching pattern that contributed to maize domestication (Studer et al. 2011). Thanks to the regulatory elements they carry, TEs have also the capacity to confer environmental responsiveness to neighboring genes (e.g. Makarevitch et al. 2015; Horváth et al. 2017; Dubin et al. 2018). Hence, the duality of TEs, seen either as purely deleterious or as powerful drivers of rapid adaptive evolution has not been resolved today and the way natural selection is acting on TEs and how they accumulate in host genomes remain important questions in evolutionary genomics (Hua-Van et al. 2011; Lisch 2013; Casacuberta and González 2013).

To achieve a more balanced view of TE evolution, one must therefore consider their accumulation as resulting from a complex balance between the rate and genomic locations at which they insert, the variety of their deleterious or beneficial effects and the rate at which they are removed from the genome through various recombination processes (reviewed in Tenaillon et al. 2010). The landscape of TE abundance across the genome provides hints about the relative impact of these different forces. In Drosophila, recombination appears to play an important role in shaping the TE landscape, as TEs are rare in regions with a high rate of recombination and their population frequency negatively correlates with recombination (Petrov et al. 2011). In contrast, TE density does not correlate with recombination in *A. thaliana* (Wright et al. 2003), but distance to the nearest gene is strongly associated with disturbance of expression (Quadrana et al. 2016). In this species, the deleterious effect of TEs thus seems be mediated directly by their presence itself rather than indirectly by their tendency to cause ectopic recombination (Wright et al. 2003). Hence, while examining abundance of TEs along a single genome provides insight into the selective forces involved, this correlative approach is inherently limited to a snapshot, with the caveat that a given pattern can arise from distinct evolutionary processes. For instance, the observation that TEs are typically found close to genes with low levels of expression can be due to either an insertion bias, a tendency of TE insertions to reduce the expression of adjacent genes, or generally weaker deleterious effects of TE insertions when adjacent genes are lowly rather than highly expressed (Maumus and Quesneville 2014; (Quadrana et al. 2016).

Different species exhibit strikingly diverse complements of TE families and superfamilies, demonstrating that evolutionary changes of this fraction of the genome can be dramatic. For instance, the majority of TEs in the pear genome (Wu et al. 2013) belong to the *Copia* superfamily, while in papaya (Ming et al. 2008) the same superfamily represents only a small fraction. It is unclear from comparing such distantly related species how fast these changes can take place, but striking differences have been observed even within species, with *e.g.* as much as 22% of genome size variation between two lines of maize mainly caused by TE differences (Vielle-Calzada et al. 2009) or a 30% increase in genome size in the Australian rice *Oryza australiensis* being caused by the recent activity of just three TE families (Piegu et al. 2006). However, because of their repetitive nature, it is generally challenging to follow the evolutionary fate of individual TE copies as soon as divergence increases. Hence, the limitation of this “global” approach is that it has limited power to pinpoint factors that prevent or promote the invasion of TEs within a given genome (Ungerer et al. 2006) (Vicient and Casacuberta 2017) (Makarevitch et al. 2015).

The Arabidopsis genus is a model of choice to study the dynamics of TEs (de Meaux and Pecinka 2012). Deep annotation by Maumus and Quesneville (2014) of the repeated fraction of the high quality genome assemblies of *A. thaliana* (Arabidopsis Genome Initiative 2000) and *A. lyrata* (Hu et al. 2011) revealed that the fraction of the genome with substantial similarity to TE sequences was more important than previously appreciated, and consisted of two distinct populations of TE sequences. Beside a large number of sequences of short, likely degraded TE-derived sequences with an ancient insertion history in both genomes, there is a massive population of recently inserted TEs in *A. lyrata* (inserted within the last million years), which is largely absent from the *A. thaliana* genome (Maumus and Quesneville (2014)). The absence of this population in *A. thaliana* has been interpreted as an effect of the shift to selfing, which occurred roughly within the same timeframe (de la Chaux et al. 2012). While theoretical models do indeed predict a major effect of the mating system on the dynamics of TEs (Charlesworth and Charlesworth 1995, Wright et al. 2008, Boutin et al. 2012), the net effect depends on the balance between sharply different processes. Therefore, it is also possible that the contrast between *A. lyrata* and *A. thaliana* is due to factors independent from the mating system. Specifically, the presence of TEs is associated with reduced levels of gene expression for TEs up to 2.5kb away in *A. thaliana,* while in *A. lyrata* TEs as close as 1kb are not associated with reduced expression of the nearby gene (Hollister et al. 2011). Furthermore, He et al. (2012) observed in F1 hybrids a consistent bias of TE transcript levels towards the *A. lyrata* copy, suggesting that *A. lyrata* TEs are less efficiently silenced than their *A. thaliana* orthologs, possibly as a result of differences in the methylation control machinery between the two species (de Meaux and Pecinka 2012). Altogether, factoring out the effect of the mating system transition is not straightforward, limiting the generality of the comparison.

To obtain a more general picture of how TEs evolve in the model Arabidopsis genus, it is essential to compare species without the confounding effect of the mating system. To follow the evolutionary fate of individual TE copies, we studied the divergence of the TE repertoires of two closely related species, *A. lyrata* and *A. halleri,* which diverged less than 1 million years ago (Roux et al. 2011) and thus remain phylogenetically close enough that TE insertions can be tracked individually. We find that both genomes host an abundant population of recently inserted TEs with almost identical insertion ages, although only a very small fraction are found at orthologous positions, indicating a very rapid turnover of these sequences. The small fraction of TE-derived sequences that is retained over the long run displays distinctive features, with gene proximity an important factor favoring TE retention. We argue that while TE accumulation in genomes has typically been studied in light of the dynamics of new insertions, their propensity for long-term retention by resisting deletion is also an important factor.

## RESULTS

### Comparing and improving genome assemblies in outcrossing Arabidopsis species

To compare TE repertoires in outcrossing Arabidopsis species, we used the high quality Sanger-based *A. lyrata* genome assembly (Hu et al. 2011),and the recently published genome assembly of the Asian subspecies *A. halleri gemmifera* (Briskine et al. 2017). To improve contiguity of the recent genome assembly of *A. halleri halleri* (Karam et al. submitted), we produced additional Illumina paired-end and mate pairs as well as PacBio sequencing reads (Table S1). We sequenced a total of (i) 12,560,731,806 base pairs using Illumina sequencing (~48x coverage of the genome) and (ii) 4,713,108,471 base pairs (~18x coverage) using PacBio sequencing with an average subread size of 3,332bp. A new Illumina-based assembly was produced combining the new reads and the reads of Karam et al. (submitted), and the PacBio long reads were used for scaffolding leading to a substantial 3-fold decrease of the number of scaffolds (9,891 to 3,152) and a 5-fold increase of the N50 (52kb to 279kb, Table1). Although the resulting assembly remains more fragmented than the *A. halleri gemmifera*, *A. lyrata* and *A. thaliana* assemblies we used in this study both in terms of the number of scaffolds and a lower N50 (Table 1), the fraction of coding sequences was roughly comparable, with only slight variations in the proportion of genic non-CDS sequences and shorter genic non-CDS sequences in *A. thaliana* (Fig. 1A), as noted previously (Hu et al. 2011). Furthermore, the higher fragmentation of the assembly affected only slightly the representation of the coding genes, since quantitative measures for the assessment of the different assemblies using Busco (Simão et al. 2015) showed similar numbers with only 46 of the 1,440 universally conserved plant genes missing from the *A. halleri halleri* assembly (3.2%) *vs.* 17 in *A. lyrata* (1.2%, Table1).

To compare TE repertoires in outcrossing Arabidopsis species, we used the high quality Sanger-based *A. lyrata* genome assembly (Hu et al. 2011),and the recently published genome assembly of the Asian subspecies *A. halleri gemmifera* (Briskine et al. 2017). To improve contiguity of the recent genome assembly of *A. halleri halleri* (Karam et al. submitted), we produced additional Illumina paired-end and mate pairs as well as PacBio sequencing reads (Table S1). We sequenced a total of (i) 12,560,731,806 base pairs using Illumina sequencing (~48x coverage of the genome) and (ii) 4,713,108,471 base pairs (~18x coverage) using PacBio sequencing with an average subread size of 3,332bp. A new Illumina-based assembly was produced combining the new reads and the reads of Karam et al. (submitted), and the PacBio long reads were used for scaffolding leading to a substantial 3-fold decrease of the number of scaffolds (9,891 to 3,152) and a 5-fold increase of the N50 (52kb to 279kb, Table1). Although the resulting assembly remains more fragmented than the *A. halleri gemmifera*, *A. lyrata* and *A. thaliana* assemblies we used in this study both in terms of the number of scaffolds and a lower N50 (Table 1), the fraction of coding sequences was roughly comparable, with only slight variations in the proportion of genic non-CDS sequences and shorter genic non-CDS sequences in *A. thaliana* (Fig. 1A), as noted previously (Hu et al. 2011). Furthermore, the higher fragmentation of the assembly affected only slightly the representation of the coding genes, since quantitative measures for the assessment of the different assemblies using Busco (Simão et al. 2015) showed similar numbers with only 46 of the 1,440 universally conserved plant genes missing from the *A. halleri halleri* assembly (3.2%) *vs.* 17 in *A. lyrata* (1.2%, Table1).

**Table 1.** Summary metrics of the four assemblies showing the relative levels of completeness and fragmentation.

|  | *A. halleri halleri* | *A. halleri gemmifera* | *A. lyrata* | *A. thaliana* |
| --- | --- | --- | --- | --- |
| Nb scaffolds | 3 152 | 2 239 | 695 | 7 |
| Total length | 174 Mb | 196 Mb | 207 Mb | 120 Mb |
| Genome cov. | 68.3 % | 76.9 % | 89.9 % | 88.9 % |
| Longest scaff. | 1.5 Mb | 4.3 Mb | 33.1 Mb | 30.4 Mb |
| N50 | 279 389 | 712 249 | 24 464 547 | 23 459 830 |
| L50 | 177 | 71 | 4 | 3 |
| CDS content | 18.7% | 19.5% | 16.3% | 35.4% |
| non-CDS gene content | 19.3% | 19.6% | 15.9% | 14.9% |
| Number of TEs | 68,583 | 85,835 | 87,477 | 39,210 |
| TE content | 15.2% | 20.8% | 27.8% | 17.4% |
| (TE content estimation) | (32.7%) | (30.2%) | (24.5%) | (19.1%) |
| Other repeats content | 4.5% | 5.4% | 5.8% | 4.3% |
| Unannotated bases | 42.2% | 34.8% | 34.7% | 28.1% |
| Complete universal single-copy orthologs | 95.3% | 97.6% | 98.5% | 98.2% |
| Fragmented universal single-copy orthologs | 1.5% | 0.3% | 0.3% | 0.5% |
| Missing universal single-copy orthologs | 3.2% | 2.1% | 1.2% | 1.3% |

**Figure 1.**
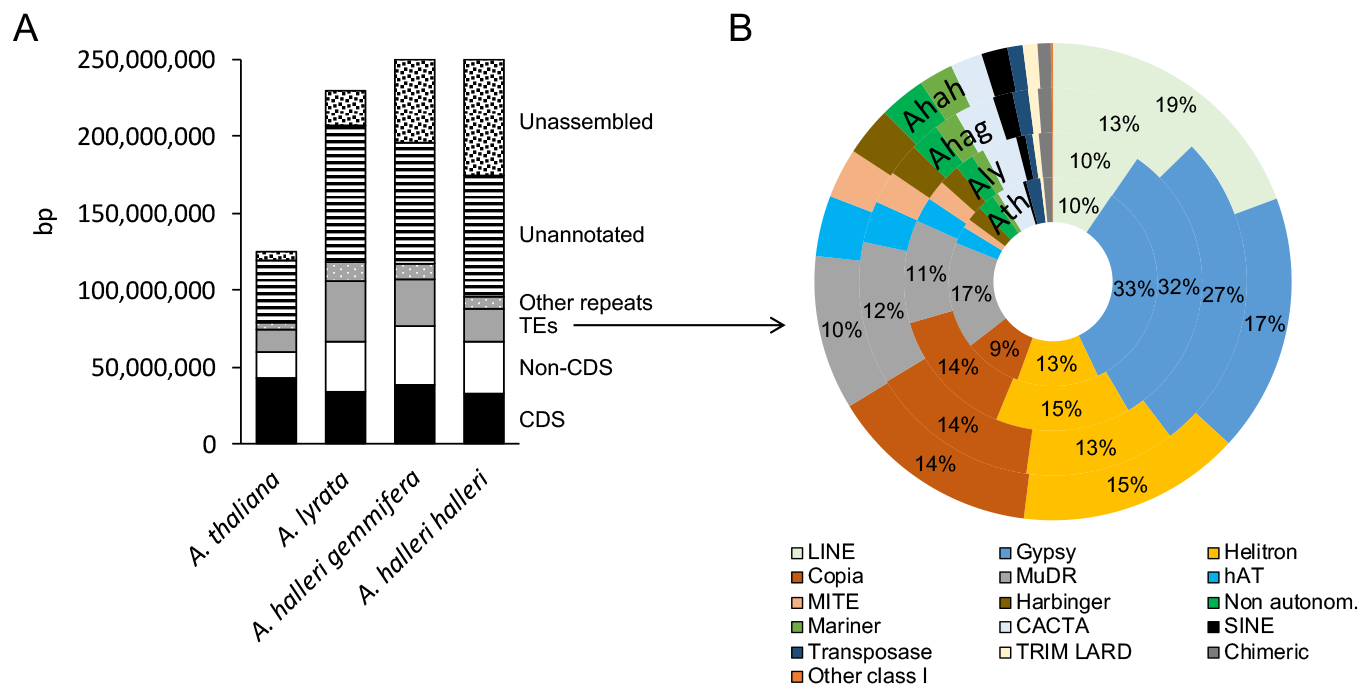
Genome composition and detailed TE content of the four assemblies A: the genomes are represented as vertical bars, split up by annotation type. For clarity, bases belonging to more than one category (overlapping annotations such as TEs included in genes) were discarded from the figure (1.84% of the total assembly size overall). Genome size estimates are from flow cytometry experiments (Johnston et al. 2005and (Briskine et al. 2017). B: Relative coverage of each TE family.

### Orthology map of genes in the assemblies

The orthology relationships between genes were defined using inParanoid (Remm et al. 2001). We identified 16,702 inparalog and ortholog clusters for *A. halleri halleri* and *A. lyrata*, in agreement with figures obtained using transcriptome data (Schvartzman et al. 2018). After removing clusters containing paralogs and applying stringent criteria (see methods), we conserved 15,620 orthologous genes between these two assemblies *i.e*. 57.5% and 47.8% of the total number of annotated genes in *A. halleri halleri* and *A. lyrata*, respectively. Reciprocal best hit Blastp approach between translated CDS of the two species led to similar results with a total of 16,900 orthologous genes (identity ≥ 85%, coverage of the query and the subject ≥ 60%). Similar numbers of inparalog and ortholog clusters were identified for human and chimpanzee using a comparable approach (Sonnhammer and Östlund, 2015). Using the same procedure, we identified 17,705 and 15,240 orthologous genes for *A. halleri gemmifera* and *A. lyrata*, and for *A. halleri halleri* and *A. halleri gemmifera*, respectively.

### Identifying and annotating TEs

To minimize bias due to annotating TEs using sequences from different reference genomes, we built libraries of consensus sequences that are representative of repetitive elements identified in each assembly separately using the *TEdenovo* pipeline of the package REPET (Maumus and Quesneville 2014). The libraries were then pooled to form a “bundle” library. Each consensus sequence was classified into types of repeats and TE superfamilies using PASTEC (Hoede et al. 2014). Finally, the bundle library was used to annotate TEs in each assembly in parallel. Overall, the bundle library was composed of 3,821 families of repeats. This library was used to annotate 68,583; 85,835; 87,477 and 39,210 TEs in *A. halleri halleri*, *A. halleri gemmifera*, *A. lyrata* and *A. thaliana*, respectively (Table 1). Our deep repeatome annotation strategy thus confirmed the higher proportion of TEs in *A. lyrata* (27.8%) than in *A. thaliana* (17.4%), as previously noted by Maumus and Quesneville 2014. Taken at face value, the proportion of TEs in *A. halleri halleri* and *A. halleri gemmifera* appears lower than in *A. lyrata* (Table 1), but this is probably not the case since these two assemblies are markedly less complete and a substantial proportion of the unassembled genome probably corresponds to repeats. To overcome this problem, we mapped the raw sequencing reads onto the bundle library, which provides an estimate of the proportion of TEs that is assembly-independent. Using this approach, we estimated that 32.7% and 30.2% of the *A. halleri halleri* and *A. halleri gemmifera* genomes are composed of TEs, with lower proportions in *A. lyrata* (25.2%) and the lowest proportion in the *A. thaliana* reference genome Col-0 (19.1%). Hence, we confirm that the three outcrosser species *A. lyrata*, *A. halleri halleri* and *A. halleri gemmifera* genomes have higher TE content than the selfer *A. thaliana*, consistent with the slightly larger genome size of *A. halleri* as compared to *A. lyrata* based on flow cytometry (Johnston et al. 2005).

Within the TE fraction, the relative proportion of the major superfamilies were roughly comparable, with Gypsy, Copia, LINE, MuDR and Helitron as the five most abundant superfamilies in all genomes, although their relative ranking varies (Fig. 1B). Hence, the higher abundance of TEs in *A. lyrata*, *A. halleri halleri* and *A. halleri gemmifera* as compared to *A. thaliana* is not due to just one TE family having expanded but rather to a more general process of accumulation over several families.

### Age distribution of TEs

The distribution of the values of identity of individual TEs to the consensus sequence of their family (classically taken as a proxy for the relative age of their insertion since TEs are initially fully identical to their copy of origin, Cordaux et al. 2010, but see Le Rouzic et al. 2013) shows that the two clearly distinct populations of TEs observed in *A. lyrata* (Maumus and Quesneville 2014) are also observed in *A. halleri halleri* and *A. halleri gemmifera*. Specifically, the distribution of percentage of identify was clearly bimodal with as many as 21,160 (30.9% of the total number of TEs) and 35,010 (40.8%) TEs with over 90% identity in *A. halleri halleri* and *A. halleri gemmifera,* respectively (Fig. 2). In the following, we define “recent” vs “old” TEs in relation to this 90% threshold. The distribution profile in *A. halleri gemmifera* is very similar to the one observed in *A. lyrata (n=32*,318 *i.e.* 36.9%) but the peak of very similar TEs is less pronounced in *A. halleri halleri*, possibly due to differences in the quality of the assembly for the most recent copies. In contrast, the number of TEs with over 90% identity above was lower in *A. thaliana* (*n*=7,938 *i.e.* only 20.2% of the total number of TEs, Fig. 2), confirming the sharply different age distribution of TEs in this species (Maumus and Quesneville 2014). Hence, the peak of putatively recent TEs observed in the *A. lyrata* genome is also observed in *A. halleri halleri* and *A. halleri gemmifera*.

**Figure 2.**
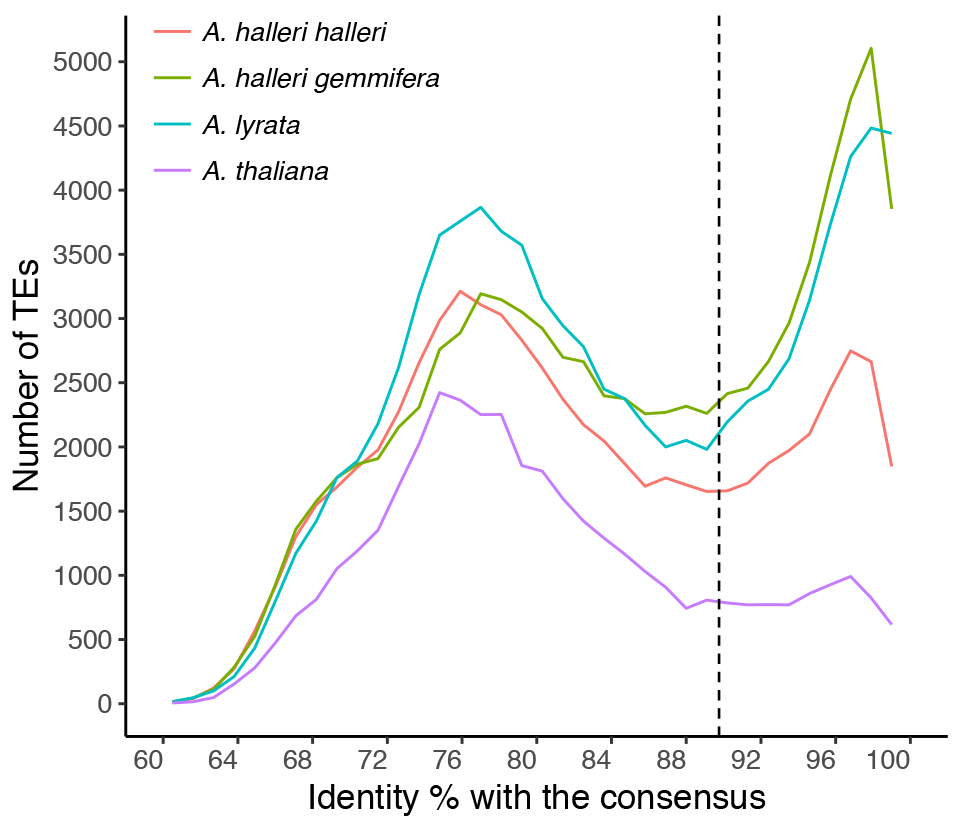
Distribution of nucleotide identity of TEs to the consensus sequence of their TE family for the three species This statistic is used as a proxy for the relative age of TE insertion. Based on this distribution, we define “old” and “young” TEs based on a threshold of 90% identity represented by the dashed vertical line (close to the lowest point of the distribution).

The different TE superfamilies differed in their contribution to the peaks of recent and ancient TEs. The LINE superfamily, for instance, had very low contribution to the recent peak, while the other four (Gypsy, Copia, MuDR and Helitron) had sometimes very sharp peaks of recent TEs (Fig. S1). Moreover, the peak of recent TEs is not caused by any single TE superfamily, but rather corresponds to the recent activity of several TE superfamilies, and hence corresponds to a general TE mobilization phenomenon. In order to evaluate the current dynamics of mobilization, we used a recently developed approach (Quadrana et al. 2016) based on the mapping of short Illumina reads from multiple individuals to detect segregating insertions that are not present in the reference assembly and show the hallmark of their recent transposition (presence of the target site duplication, TSD). We found that the superfamily composition of this set of presumably currently active copies in 54 *A. halleri gemmifera* individuals (Kubota et al. 2015) is very similar to that of the peak of recent TEs present in the assembly (Fig. 3 and S1). Hence, TE mobilization appears to be ongoing and the relative contribution of the different superfamilies seems to have remained relatively stable in the recent past.

**Figure 3.**
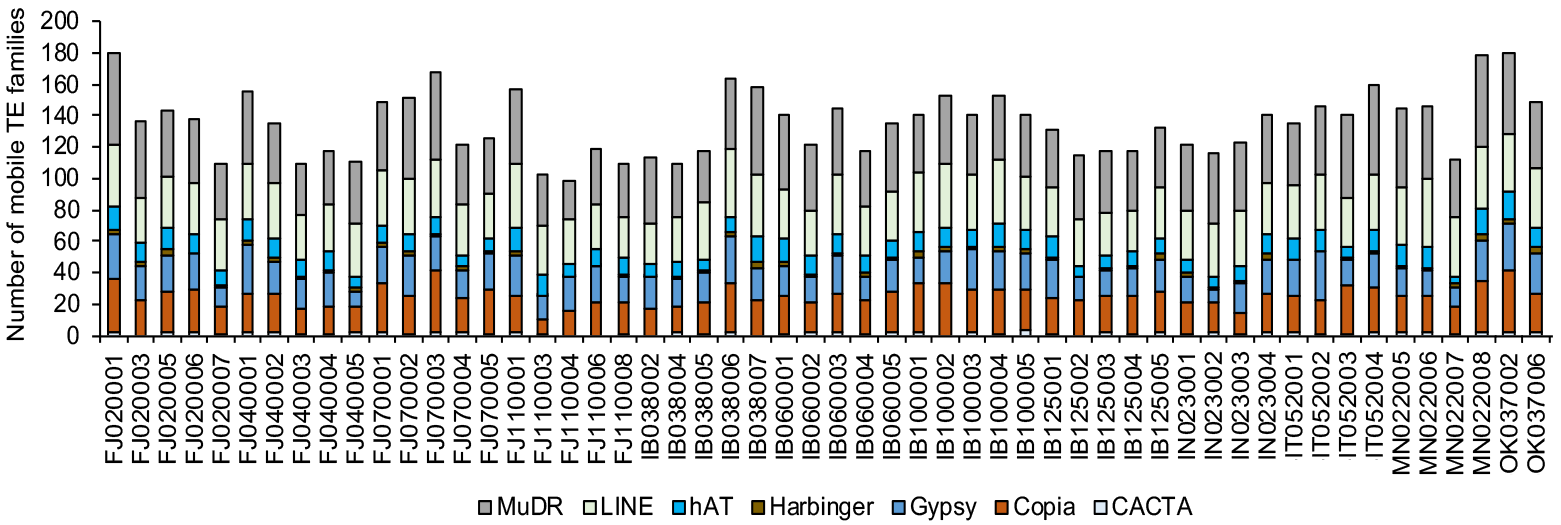
Mobilome composition and variation among *A. halleri gemmifera* accessions

### Low proportion of orthologous TEs in spite of recent divergence

Next, we sought to follow the fate of individual TE copies between pairs of lineages in order to distinguish TEs that have been either specifically inserted or deleted in one of the two lineages from TEs that have been maintained at orthologous positions since the divergence between pairs. To define TE orthology in a context where the compared assemblies exhibit different levels of contiguity, we used stringent positional information determined by the identity of the pair of flanking genes with a strict one-to-one orthology relationship between the two genomes compared. The presence of a TE in the orthologous intergenic interval was then determined based on a relaxed Blast search procedure. For TEs within genic sequences, we searched for the presence of a TE in the unambiguous ortholog, when it existed. In turn, to avoid spurious results due to multiple hits that may arise because of the relaxed Blast criteria, we restricted the analysis to orthologous intergenic segments shorter than 70kb and discarded TEs that are either on contigs with no orthologous gene or on the extremity of contigs. Using this set of conditions, the intergenic segments considered contained an average of 2.7 distinct TE sequences, thus enabling us to cross-check their presence (orthology) and absence (non-orthology) in the two genomes with substantial accuracy. In spite of the use of relaxed Blast parameters, our analysis identified only 8,394 orthologous TEs between *A. halleri halleri* and *A. lyrata*, representing a minority of the TEs. Specifically, this number of orthologous TEs represents 28.8% of the 29,111 TEs in *A. halleri* and 21.6% of the 38,919 interrogated TEs from *A. lyrata* (Fig. 4A). As expected, a higher proportion of orthologous TEs was detected when comparing the very closely related *A. halleri halleri* and *A. halleri gemmifera (i.e.* at the sub-species level, 51.6% and 38.3%; Fig. 4B).

**Figure 4.**
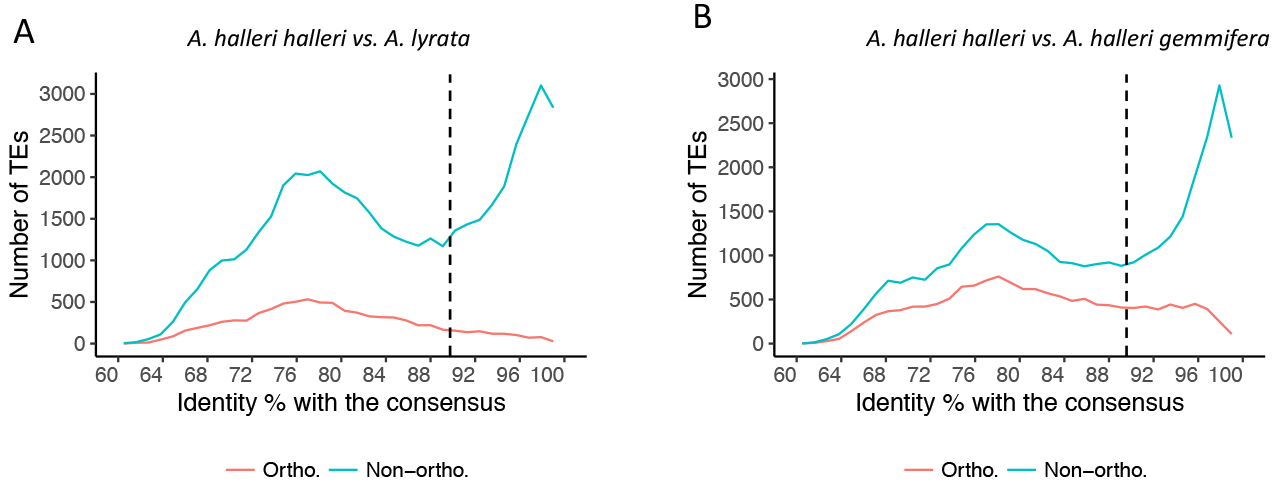
Distribution of nucleotide identity of TEs to the consensus sequence of their TE family

The age distribution of orthologous TEs (as defined by the divergence to their consensus) was strikingly different from that of non-orthologous TEs. As expected, orthologous TEs were almost exclusively old with a very small fraction belonging to the population of recently inserted TEs, whereas non-orthologous TEs were either anciently or recently inserted, with a relative proportion of these two categories closely matching that of the overall genome (Fig. 4, Table 2). Similar results were observed in the comparison between *A. halleri gemmifera* and *A. lyrata* (Table S2). Given the time scales considered, recently inserted TEs may either have inserted after the species became isolated, or have been present in the ancestor some time before the split and have been removed in one of the two species. It is therefore impossible to unambiguously reconstruct their evolutionary history.

**Table 2.** Proportion of orthologous and non-orthologous TEs in pairwise comparisons.

| Pairwise comparison | *A. halleri halleri* vs *A. lyrata* | *A. halleri halleri* vs *A. halleri gemmifera* |
| --- | --- | --- |
| Total orthologous TEs | 8,394 | 14,720 |
| Old orthologous TEs | 7,471 | 11,536 |
| Young orthologous TEs | 923 | 3,184 |
| Total non-orthologous TEs | 51,242 | 37,534 |
| Old non-orthologous TEs | 31,471 | 21,270 |
| Young non-orthologous TEs | 19,771 | 16,264 |

### Factors associated with long-term maintenance of ancient TEs

In contrast, reconstructing the evolutionary history of ancient TEs (<90% sequence identity to their respective consensus, Fig 4) is relatively straightforward. We note that, by definition, this population of TE-derived sequences has accumulated mutations since their insertion, and so corresponds to largely degraded and likely inactive copies rather than full-length elements. Assuming identical rate of divergence, “old” TEs had to be present in the ancestor species, so that their absence in one genome can be readily interpreted as resulting from a deletion process. Based on this assumption, we sought to identify the factors associated with long-term maintenance of individual TEs. First, we compared the different superfamilies and found that old members from the Helitron superfamily were preferentially maintained in the long-term relative to others, since this class of TEs was more represented among orthologous (25.6%) than among non-orthologous TEs (11.6%, Fig. 5A). Conversely, old members of the LINE and Copia superfamilies were enriched in the non-orthologous fraction (26.4% and 18.9% for LINE and Copia, respectively) relative to the orthologous fraction (11.3 and 9.2%, respectively) and were therefore more rapidly deleted. Second, we found that TEs that had been maintained at orthologous positions tend to be on average 19.8% shorter than those that had been deleted from one of the two genomes (322.8 *vs.* 401.6 bp, *p*<2.2e^−16^), but comparing the medians of the two distribution showed the opposite pattern, suggesting that the difference is largely driven by a limited set of large TEs that are only found in the non-orthologous fraction (Fig. 5B). Third, we compared the location of orthologous *vs.* non-orthologous old TEs and observed that orthologous TEs tended to be found more often within genes than non-orthologous TEs (Fig. 5C). For instance, in the *A. lyrata vs*. *A. halleri halleri* comparison, 30.3% of orthologous TEs but only 15.8% of non-orthologous TEs were found within genic sequences (*p*<2.2e^−16^). The same qualitative pattern was true for the *A. lyrata* vs *A. halleri gemmifera* comparison (Fig. S2) and the *A. halleri halleri vs. A. halleri gemmifera* comparison, albeit with a lesser contrast for the latter (22.2 *vs.* 19.8%, Fig. 6C). For old TEs within genic sequences, we further distinguished between TEs within CDS and non-CDS sequences. We observed that old TEs located within CDS are more likely to be retained at the orthologous state than TEs located in non-CDS sequences (Fig. 5D). In fact, around 35.5% of orthologous TEs were found within CDS sequences, while they were only 11.8% for non-orthologous TEs (*p*<2.2e^−16^). In line with this observation, we also observed that TEs outside genic regions tended to be retained more readily when located close to genes (Fig. 5E). Among the old TEs, those that have been retained at orthologous positions between *A. halleri halleri* and *A. lyrata* were located on average 1,829 bp away from their closest gene, while those that have been retained either in *A. halleri halleri* or in *A. lyrata* only (and thus have been deleted from the other lineage) were located on average at a distance of 2,303 bp, *i.e.* they were located 26% farther (*p*<2.2e^−16^). Overall, these results suggest that TEs in gene-rich regions tend to be protected from deletion, possibly because of the deleterious effects associatted with the imprecise nature of the deletion process, which tend to remove flanking sequences as well. We then sequenced small RNAs from the *A. lyrata* MN47 accession and compared the proportion of old TEs with substantial siRNA production (>5 uniquely mapped reads per million reads and covering at least 10% of the length of the TE) as a proxy for efficient targeting by the RdDM pathway. As explained above, recent TEs were discarded from this analysis because of their ambiguous evolutionary history. Old TEs located within genes were less often targeted by the RdDM pathway than those outside of genes (*p*<2.2e^−16^) (Fig. 5F). For TEs located within genes, we found that old TEs that have remained orthologous were less likely to be RdDM targets than those that have been deleted since divergence, with 10.3% of non-orthologous TEs showing active siRNA production *vs.* only 2.3% for orthologous TEs (*p*<2.2e^−16^). However, for TEs located outside of genes we found no difference in siRNA production by orthologous vs. non-orthologous TEs (17.6% and 17.5% respectively, *p*=0.9315). Since the mechanism of TE silencing operates through DNA methylation, we further compared the level of methylation of orthologous and non-orthologous TEs in *A. lyrata,* taking advantage of the bisulfite DNA methylation data available for *A. lyrata* (Seymour et al. 2014). In accordance with the siRNA mapping analysis, we found that orthologous TEs within genes were less methylated than any other populations of TEs (*p*<2.2e^−16^, Fig. 5G). Indeed, they presented a mean percentage of methylation of only 4.0% compared to 22.8, 20.3, 32.4% for non-orthologous TEs within genes, orthologous TEs outside of genes, and non-orthologous TEs outside of genes, respectively. These results suggest that a low siRNA production and low DNA methylation levels are associated with the long-term maintenance of old TEs within genes. In contrast, these two factors may not be related to the long-term maintenance of TEs outside of genes.

**Figure 5.**
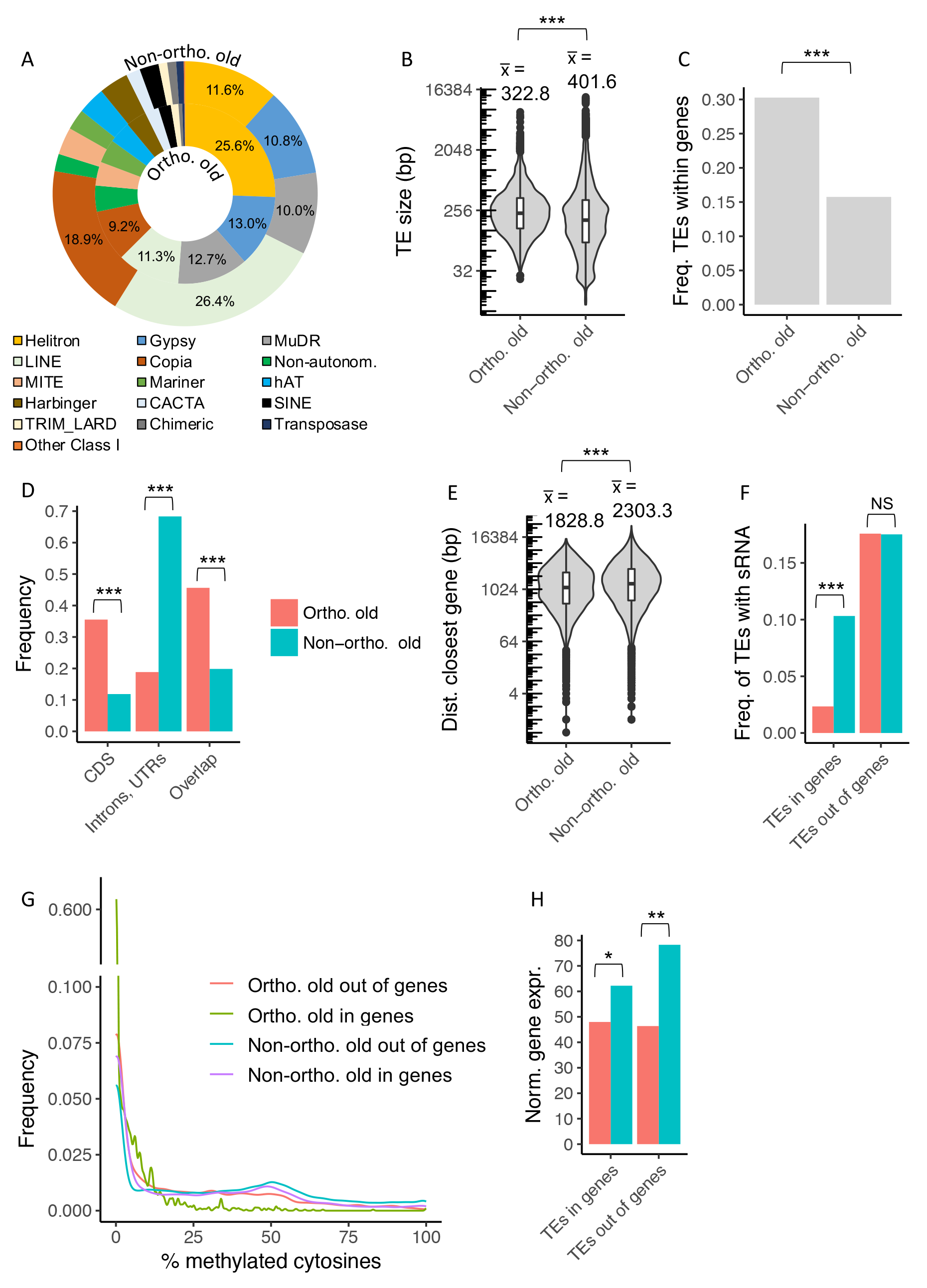
Identification of factors related to the long-term maintenance of old TEs using the comparison between *A. halleri halleri* and *A. lyrata* A: superfamily composition, B: TE length, C: frequency of orthologous and non-orthologous TEs within genic sequences, D: frequency of orthologous and non-orthologous TEs within different categories of genic sequences, E: distance to the nearest gene for TEs outside of genes, F: frequency of TEs with active siRNA production, as defined by the presence of at least 5 reads and siRNA reads covering at least 10% of the total length of the TE sequence G: frequency of TEs related to the percentage of methylated cytosines, H: normalized gene expression for genes containing a TE or genes without a TE. Statistical significance is indicated using the following code: ^“***”^ for *p<*0.001, ^“**”^ for *p* between 0.01 and 0.01, ^“*”^ for *p* between 0.01 and 0.05, ^“.”^ for *p* between 0.05 and 0.1 and “NS” for P>0.1.

**Figure 6.**
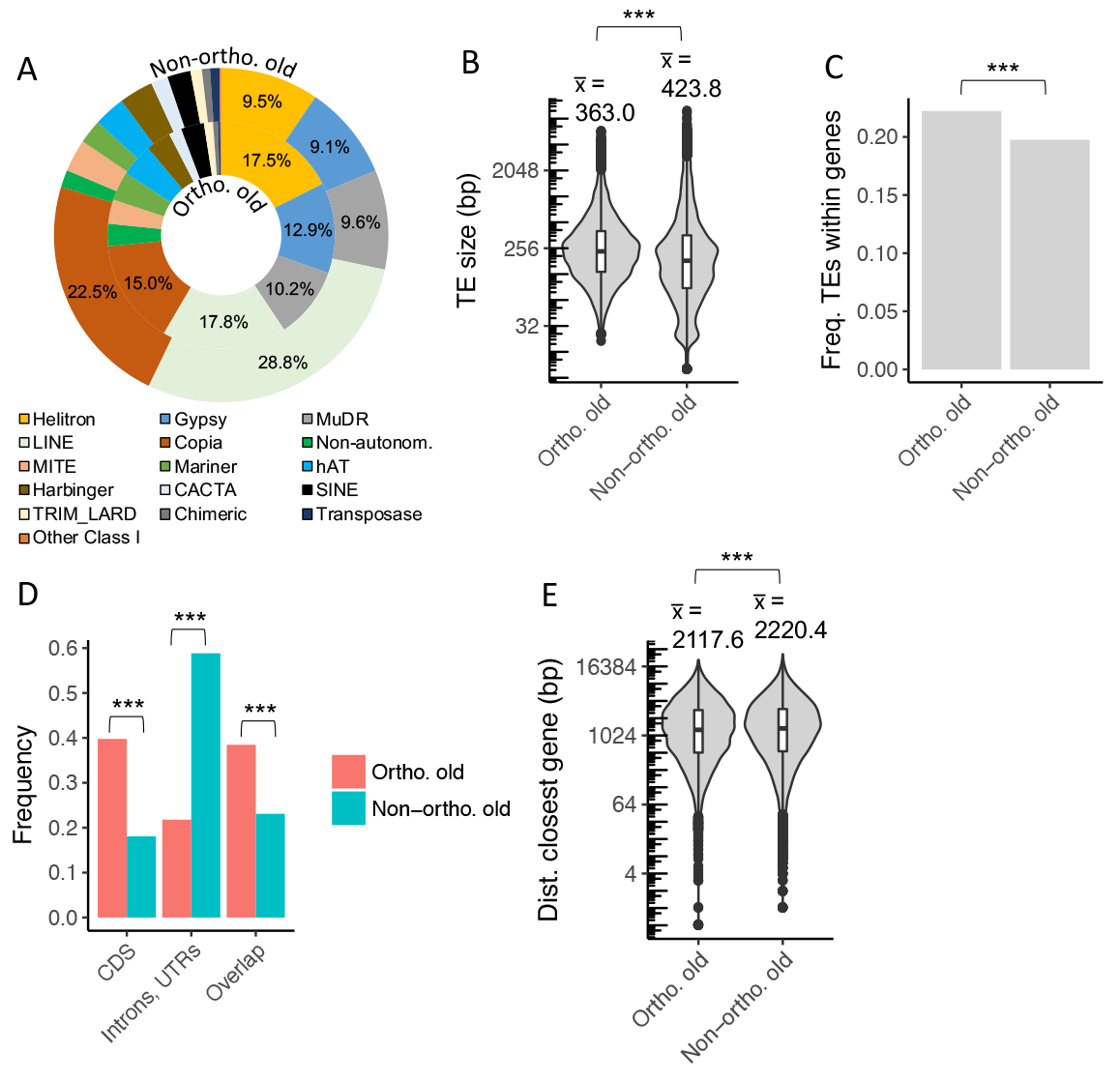
Identification of factors related to the long-term maintenance of TEs using the comparison of the TE content between *A. halleri halleri* and *A. halleri gemmifera* A: superfamily composition, B: TE length, C: frequency of orthologous and non-orthologous TEs within genic sequences, D: frequency of orthologous and non-orthologous TEs within different categories of genic sequences, E: distance to the nearest gene for TEs outside of genes. Statistical significance is indicated using the following code: ^“***”^ for *p<*0.001, ^“**”^ for *p* between 0.01 and 0.01, ^“*”^ for *p* between 0.01 and 0.05, ^“.”^ for *p* between 0.05 and 0.1 and “NS” for P>0.1.

Finally, we used RNA-seq data from the same *A. halleri halleri* accession to compare the expression of genes containing old TE sequences that have been either retained or removed since the separation of the two species. We reasoned that if TEs are deleterious on average, removing them should be advantageous even in the face of the deleterious effect of local deletions. If so, TE removal should occur more readily close to or within genes with high expression than genes with low expression. Accordingly, we found that on average genes with an orthologously-maintained TE in their DNA sequence were expressed at slightly lower levels than the genes with a TE that has been removed from *A. lyrata* (*p=*0.01085, Fig. 5H). The same pattern was also found for TEs in intergenic regions when comparing for each TE the expression of the closest gene along the chromosome (*p=*0.00175).

## DISCUSSION

### Dynamics of TE accumulation in two outcrossing species

Overall, the population of TEs in the two *A. halleri* assemblies that we studied is very similar to the one described in *A. lyrata*, both in terms of TE families present and in their age distribution. As noted previously, these profiles are sharply distinct from that seen in *A. thaliana* (Hu et al. 2011; de la Chaux et al. 2012; Slotte et al. 2013; Maumus and Quesneville 2014). This contrast has been attributed either to differences in the mating system (Wright and Schoen 1999; Boutin et al. 2012) or to a specific burst along the outcrosser lineages (He et al. (2012); de Meaux and Pecinka 2012). Although our analysis cannot formally distinguish between these two possibilities at this stage, our results unambiguously demonstrate that the dynamics of TE accumulation that is shared between *A. halleri* and *A. lyrata* has been in place at least since their divergence, *ca.* 1 Myrs ago and is therefore not an event of the very recent past.

This dynamic is first characterized by the fact that multiple TE families are currently active. The young population of TEs observed in *A. halleri* and *A. lyrata* is composed of several families, with Helitron, Gypsy, MudR and LINE being the most contributing families. This is confirmed by the analysis of segregating (neo)insertions that are absent from the reference assembly in *A. halleri gemmifera*. Overall this pattern is similar to what was observed in the *A. thaliana* genome (Quadrana et al. 2016), albeit to an even greater scale and comes in stark contrast to the human genome, where only a few TE families contain mobile copies, all belonging to the LINE-1 and SINE families (Richardson et al. 2015). In spite of the recent divergence between the two species we find very few orthologous TEs between *A. halleri* and *A. lyrata*, even for the population of “older” TE-related sequences that must have been present before speciation. Overall, even though the dynamics of TE accumulation seems to be shared, the resulting TE fractions of the two genomes are very different, indicating a rapid turnover of TE-related sequences. It will now be essential to compare quantitatively the rate at which TEs transpose and get removed between different species and how these rates are affected by various biological features. TEs have been used as phylogenetic markers in other taxa (e.g. birds; Han et al. 2011), where the rate of turnover of TE-related sequences seems to be slower. The rate of DNA loss varies extensively across species (Petrov et al. 2000), but the determinants of this variation are poorly understood. Whether the rate and pattern of TE removal differ from the more general process of non-coding DNA loss across the genome is an important question for the future.

### Factors associated with TE deletion or maintenance

Given the very rapid elimination of old TEs that we observe, how can a substantial number of old TEs be maintained for a long period of time, while a complete elimination would have been expected if this was a continuous process? We found marked differences in the propensity of TE-related sequences to resist deletion and therefore be maintained over a time scale of *ca.* one million years of total divergence.

### Long-term maintenance of helitrons, rapid removal of LINE and Copia

First, our analyses suggest that Helitron elements are more likely to maintained over the long-term in *A. halleri* and *A. lyrata*. As noted by Maumus and Quesneville 2014, helitrons tend to have lower GC content than the other TE superfamilies, which may be associated with reduced targeting by RDdM because less cytosines are available for methylation, hence leading to less disruption of neighbouring gene expression. It is tempting to speculate that the lower GC content of helitrons may make them less deleterious, allowing for their preferential long-term maintenance. In contrast, LINE and Copia families are those that have been the most strongly eliminated since the divergence of the two species. Mao and Wang (2017) recently observed that in grass, SINE families were retained over the long term. Like in grass, SINEs have low abundance in the *A. halleri* and *A. lyrata* genomes, since they cover less than 3% of the repeat sequences. However, in these species they do not seem to be associated with a long-term maintenance, as they are equally represented amongst old TEs that have been maintained at orthologous positions and amongst those that have been deleted from one of the two genomes (Fig. 4). Hence, the long-term maintenance of particular TE families seems to be lineage-specific and cannot easily be generalized.

### Sheltering of TEs by proximity to genes

A striking observation is the long-term retention of TE sequences in gene-rich regions. As we focused on the population of “old” TEs that had to be present in the most recent common ancestor, this pattern is unlikely to be caused by an insertion bias of recent specific insertions towards genic regions and rather reflects a process of differential retention. Alternatively, this pattern may also be caused by gene-rich regions being better assembled, resulting in TEs in those regions being more readily found in the different assemblies. This effect is likely minor because our analysis focuses on old TEs, which should be relatively less problematic in terms of assembly because they tend to be less identical across copies. Also, in this case the most poorly assembled genomes should show less non-orthologous TEs, while here the reciprocal analyses provide similar results. The *A. lyrata* genome sequence is a high quality assembly obtained using the Sanger technology, yet does not show a specifically elevated fraction of non-orthologous TEs. Clearly, long-read technologies should resolve this issue (Chakraborty et al. 2018).

The relative enrichment of orthologous TEs in genic sequences as compared to non-orthologous TEs is consistent with the interpretation that TEs within gene-rich regions benefit from a “sheltering” effect, whereby a deletion of the TE sequence involves the risk of also deleting part of the gene sequence, which would be highly deleterious in particular when they have become integrated within coding sequences. Hence, the effective rate of deletion might be higher for non-genic TEs than for genic TEs, resulting in a long-term enrichment of the sheltered genic TEs. This process of differential retention was less pronounced when comparing the more closely related *A. halleri halleri* vs. *A. halleri gemmifera* species, indicating that such differential enrichment is a relatively slow process. In grass, Mao and Wang (2017) showed that members of the SINE TE family are often shared across species and are also enriched in and near protein coding genes, possibly as a result of differential removal of SINE copies in gene-poor regions.

### TE deletion: a cure worse than the disease?

Our results suggest that several factors can affect the long-term retention of transposable element sequences, and in particular the proximity to highly expressed genes. We propose that the process of differential TE retention results from the balance between the deleterious effects of the TE itself and that of the deletion removing it. While the presence of TE sequences was shown to equally frequently increase or decrease gene expression (Stuart et al. 2016) or to have no direct causal effect (Maumus and Quesneville 2014) in *A. thaliana* (but see Uzunović et al. 2018), we found that the more highly expressed genes rarely retain orthologous TEs. This suggests that selection in favor of TE deletions varies according to the level of gene expression, with deleterious effects of TE presence generally outweighing the cost of their eventual deletion when they are close to highly expressed genes.

Earlier studies have shown that the rate of DNA loss can be highly heterogeneous across genomes (Petrov et al. 2000). It is possible that the level of sequence identity among repeated sequences may contribute to this variation, as more identical sequences are more likely to be involved in the heterologous recombination that is believed to be responsible for DNA deletions. If so, the most recently inserted TEs would be expected to show an even faster elimination, as proposed by Maumus and Quesneville (2016). This might also contribute to decrease the proportion of young orthologous TEs. Beside the fact that they might have been inserted after the species divergence (but as we explained, precisely dating these events is challenging), they might be eliminated even more rapidly than old TEs that recombine less easily. Hu et al. (2011) suggested that the *A. thaliana* genome is characterized by ongoing positive selection on deletions, favoring genome shrinkage (but see Long et al. 2013). It would be interesting to determine how many of these deletions involve the removal of TE sequences.

In addition to this “sheltering” effect of TEs considered as deleterious or quasi neutral elements, it is also possible that those TEs that are retained in the long term have acquired a functional beneficial role for their host genome (being “domesticated”), thus making their removal deleterious in itself. It is unclear how frequent this phenomenon might be, but several clear examples of such domestication have been reported in the literature, including the regulation of stress-response genes by acquisition of response elements carried by some TEs (Capy et al. 2000; Makarevitch et al. 2015) or the production of siRNAs that trigger the trans-silencing of active relatives and therefore contribute to immune memory (Lisch 2009; Bousios and Gaut 2016; Roessler et al. 2018). A recent study however showed that TE exaptation for regulatory function is rare, and is mostly associated with “old” TEs, suggesting a model in which TE-derived sequences are initially repressed, after which a small fraction acquires and retains enhancer activity (Simonti et al. 2017). Clearly, among the repeat sequences, the old orthologous TEs that we identified here are the most likely to have acquired advantageous biological functions. Better understanding the variety of factors causing differences in retention propensity will now be an exciting and interesting next step.

## MATERIAL AND METHODS

### A. halleri genome de novo assembly

Assembling genomes of outcrossing organisms is a challenging task because outcrossing involves a high level of heterozygosity. To increase contiguity of the recently released *A. halleri halleri* assembly based on Illumina reads (Karam et al. submitted), one paired-end (PE) and two mate-pair (MP) additional libraries were prepared from the same accession PL22-1A with the TruSeq PCR-free and the Nextera DNA library prep kits (Illumina, California, United States), respectively (Table S1). We additionally produced PACBIO sequences (6 SMRT cells). Quality of the Illumina raw reads was assessed using FastQC (version 0.10.1, http://www.bioinformatics.babraham.ac.uk/projects/fastqc/) and reads were filtered accordingly using Trimmomatic (version 3, Bolger et al. 2014). When present, Ns were removed using prinSeq (version 0.20.4, Schmieder and Edwards 2011). The total number of filtered Illumina reads (Karam et al. submitted and this study) represented a 110x coverage of the *A. halleri* estimated genome size (~ 255 Mbp, Johnston et al. 2005). A new *de novo* assembly was carried out with the AllPathsLG assembler (version r44837, Gnerre et al. 2011) using all PE and MP reads. The kmer spectrum analysis carried out by AllPathsLG estimated the *A. halleri* genome size to 266 Mb, with ploidy equal to 2 and a SNP rate of 1/150, consistent with previous estimation (Karam et al. submitted). This initial AllPathsLG assembly was then improved with the following strategy: (i) a scaffolding step was performed using the PACBIO reads and followed by a gap filling step using the PE and MP Illumina reads and (ii) a second step of scaffolding was performed using the PACBIO reads. Scaffolding was carried out using the SSPACE-LongRead.pl perl script of SSPACE (version 1.1, Boetzer and Pirovano 2014) and gap closing was achieved using the GapFiller.pl perl script of gapfiller (version 1.10, Nadalin et al. 2012). Gene annotation was based on Maker (version 2.31.8, Campbell et al. 2014). EST evidence, protein homology and repeat masking references were provided from *A. thaliana*. Gene prediction was allowed from EST inference and from protein homology and resulted in the prediction of 27.992 genes. Genome metrics were obtained using QUAST (version 4.0, Gurevich et al. 2013) and genome assembly and annotation completeness was assessed with BUSCO (version 3, Simão et al., 2015) using the Embryphyta odb9 dataset composed of 1,440 universal single-copy orthologs.

### TE annotation

In order to produce a genome-wide annotation of repetitive sequences, the four genomes were annotated using the package *REPET* (version 2.5, Flutre et al. 2011), which is composed of two main pipelines, dedicated to *de novo* detection, annotation and analysis of repeats, in particular TEs, in genomic sequences (Fig. S3). Briefly, the first pipeline, *TE de novo*, starts by comparing the genome with itself and clusters matches. Then, for each cluster, it builds a multiple alignment from which a consensus sequence is obtained. Finally, consensus sequences are classified according to TE features, and redundancy is removed. The second pipeline dedicated to the annotation (*TE annot*) involves several steps, including TE detection by similarity search, spurious hit removal and connection of distant fragments. In our study, a library of classified, non-redundant consensus sequences was obtained by combining the *TE de novo* analysis performed on the four species. Then, the bundle library was used to annotate each of the four genomes separately using *TE annot*.

In parallel, the proportion of TEs in each of the four genomes was estimated using an assembly-free approach. The raw sequencing reads that mapped onto the bundle library using Bowtie2 were considered as representing TEs and the other reads as non-TEs sequences. The genomic Illumina reads were obtained from Karam et al. (submitted, 37,262,746 reads in total) for *A. halleri halleri*, or downloaded from the NCBI SRA database: DRR013376 (38,782,027 reads) for *A. gemmifera*, SRR2040788 and SRR2040789 (48,602,962 reads in total) for *A. lyrata* and ERR1399719 (38,425,727 reads) for *A. thaliana*.

### TE orthology

TE orthology relationships were obtained for each pairwise comparison *i.e. A. halleri halleri vs. A. lyrata*, *A. halleri halleri vs. A. halleri gemmifera* and *A. halleri gemmifera vs. A. lyrata* using the orthology of genes as detailed in Fig. S4. Briefly, an orthology map of genes, using CDS annotations (excluding all CDS annotations included in TE annotations), was constructed with Inparanoid (Remm et al. 2001). The *A. thaliana* genome was used as outgroup in the comparison of *A. halleri halleri* or *A. halleri gemmifera vs. A. lyrata*, whereas the *A. lyrata* genome was used as outgroup when comparing *A. halleri halleri* and *A. halleri gemmifera*. To avoid spurious hits, a stringent score cut-off of 100 bits was applied, paralogs were eliminated from the analysis, and only clusters with bootstrap values ≥99% for each of the two orthologs were conserved. Then, we selected only TEs located between two genes of this orthology map (called “framed” TEs, or TEs located within a genic sequence (“inserted” TEs). For each “framed” TE in one species, a blast search was performed between the TE sequence and the genomic sequence between the same pair of orthologous genes in the other species. We restricted this analysis to chromosomal segments of at most 70 kb (from either the subject or the query genome). Similarly, for “inserted” TEs, the TE sequence was compared with the orthologous gene sequence. Both “framed” and “inserted” TEs presenting a blast hit with an E-value ≤1E^−10^, an identity ≥ 80% and at least some overlap with a TE annotation were defined as orthologous. The other TEs, which did not fulfill these three criteria, were defined as non-orthologous. These criteria correspond to a relatively relaxed search and should result in a strong power to detect orthologous TEs, resulting in a conservative analysis.

### TE analyses

TEs were separated into “young” and “old” classes according to whether they reached the cut-off of 90% identities with their cluster consensus sequence, or not. Differences in the size and distance to nearest gene were tested using a non-parametric Mann-Withney test. Differences in the proportion of genic *vs.* non-genic TEs and in the proportions of CDS vs non-CDS TEs were tested using a χ2 test with 1 and 2 degrees of freedom, respectively.

### Identification of segregating non-reference (neo)insertions

We used a modified version of the pipeline developed by Quadrana et al. (2016) to identify segregating non-reference (neo)insertions in the large population sample of 54 *A. halleri gemmifera* individuals (SRA #DRA003268, omitting samples OK037001 and OK037003 because of low coverage). Basically, this pipeline was modified to consider both discordant and split-reads to call insertions. The analysis has two steps. We first performed *de novo* detection of non-reference TE insertions, for which put a threshold of at least ten supporting reads (discordant-reads + split-reads). We then assessed the presence or absence of these putative non-reference TE insertions across the whole population by relaxing the parameters (at least two discordant-read and/or split-read) used to detect them in the first place. This improved the discovery of putative TE-insertions that are shared by more than one accession.

### siRNA mapping and DNA methylation analyses

Total RNA was isolated from leaves of *A. lyrata* MN47 using the Qiagen miRNeasy Mini Kit (catalog # 217004). A total of 3 μg of RNA was sent to LC sciences (Houston, TX, USA) were an Illumina TruSeq Small RNA library was constructed and sequenced, leading to the obtention of approximately 14 million 1×50 bp reads. Adaptators were removed from the Illumina reads using Cutadapt (version 1.2.1, Martin 2011) and reads were cleaned using Prinseq (version 0.20.4, Schmieder and Edwards 2011) with specified parameters:-min_len 15 –max length 25 – noniupac-min_qual_mean 25-trim_qual_right 20-ns_max_n 0. The quality of the Illumina cleaned reads was checked using FastQC (version 0.11.4, https://www.bioinformatics.babraham.ac.uk/projects/fastqc/). The sequences were deposited in the NCBI-SRA database (accession no. XXX). rRNAs, tRNAs, snRNAs and snoRNAs were removed from the sRNAs sequences through Bowtie (version 1.0.0, Langmead et al. 2009) alignments using a set of 7,743 eukaryotic sequences obtained from NCBI database corresponding to these types of non-coding RNAs. sRNA reads were mapped on the *A. lyrata* MN47 genome (Hu et al. 2011) using Bowtie. Multiply mapping reads were discarded and only alignments presenting no mismatch were conserved. A TE was defined as producing substantial siRNAs when it presented an overlap with more than 5 reads per million and when it was covered on more than 10% of its length.

The DNA methylation matrix for *A. lyrata* MN47 of Seymour et al. (2014) was used to evaluate the methylation status of orthologous vs non-orthologous TEs. Following Seymour et al (2014), we considered a cytosine site as significantly methylated when its methylation rate was ≥20% in at least one of the four tissue-treatment combinations (shoot, root, 4°C, 23°C). Then, we calculated for each TE the percentage of methylated sites.

### Gene expression analysis

To evaluate gene expression, we generated RNA-seq data from shoot of *A. halleri* PL22-1A plants cultivated in standard greenhouse conditions. The number of reads mapped on each transcript of the PL22 reference transcriptome (Schvartzman et al. 2018) were counted and normalized (TPM) (Li and Dewey 2011). Correspondence between transcripts from the reference transcriptome and gene models in the assembly was established by BLAST using a stringent criteria (95% identity over at least 100 bp).

## ACKNOWLEDGEMENTS

We thank Juliette de Meaux, Arnaud Le Rouzic, Marie Mirouze, Maud Tenaillon, Anna-Sophie Fiston-Lavier and Xavier Vekemans for insightful discussions. This work was funded by the European Research Council (NOVEL project, grant #648321), by Région Hauts-de-France (MICRO2 project, Program “projet emergent”), by the Fonds de la Recherche Scientifique–FNRS (PDR-T.0206.13) and by the University of Liège (SFRD-12/03). The ANR ELOCANTH (grant ANR-12-JSV7-0010) provided resources for plant culture. The authors also thank the Région Hauts-de-France, the Ministère de l’Enseignement Supérieur et de la Recherche (CPER Climibio) and the European Fund for Regional Economic Development for their financial support. M.H. is Research Associate of the FNRS.

---

**Figure S1.** Distribution of identity of TEs to the consensus sequence of their TE family, separated by superfamily

For each species, superfamilies are sorted according their contribution to the peaks of the most recent population of TEs (using a threshold of 98%).

**Figure S2.** Identification of factors related to the long-term maintenance of TEs using the comparison of the TE content from *A. halleri gemmifera* and *A. lyrata.* A: distribution of nucleotide identity of TEs to the consensus sequence of their TE family, B: superfamily composition, C: TE length, D: frequency of orthologous and non-orthologous TEs within genic sequences, E: frequency of orthologous and non-orthologous TEs within different categories of genic sequences, F: distance to the nearest gene for TEs outside of genes. Statistical significance is indicated using the following code: ^“***”^ for *p<*0.001, ^“**”^ for *p* between 0.01 and 0.01, ^“*”^ for *p* between 0.01 and 0.05, ^“.”^ for *p* between 0.05 and 0.1 and “NS” for P>0.1.

**Figure S3.** Pipeline used for the deep repeatome annotation.

**Figure S4.** Strategy to identify orthology relationships of TE sequences as determined by positional information from the flanking genes

The first step consists in defining the orthology of genes (blue squares) between genomes X and Y using Inparanoid. In our example, A/A’, C/C’ and E/E’ are considered as orthologous pair of genes (represented by blue arrows). The orthology of TEs is defined sequentially for genome X and Y but the process are similar: only TEs between two orthologous genes spaced for at most 70kb (black squares named a and b in our example) (named “Framed”) and TEs located within genes (black square c) (named “Inserted”) are analysed. TEs which are located at an extremity of a scaffold (d and d’) and TEs located on scaffold without orthologous genes are discarded. The sequence of the TE a and b, which are located between A and C genes of the orthology map are compared using Blastn (thresholds: Evalue ≤1E^−10^, an identity ≥ 80%) to the sequence between the orthologous genes of A and C, *i.e.* A’ and C’. The TE a presents a blast hit, and a TE annotation overlaps the Blast hit in genome Y. Hence a and a’ are defined as orthologous. No-significant blast hit is retrieved for b, which is defined as non-orthologous. The sequence of the TE c located within the E gene is compared to the sequence of the E’ gene. In our example, we considered that the Blast hit is significant and overlaps a TE annotation in Genome Y. The TE c is defined as orthologous.

**Table S1.** Summary statistics of input sequence data for *de novo* assembly of the *A. halleri* genome. Libraries marked with an asterisk were produced and sequenced by (Karam et al., submitted).

**Table S2.** Proportion of orthologous and non-orthologous TEs in *A. halleri gemmifera* and *A. lyrata* genomes

